# Polymer microparticles prolong delivery of the 15-PGDH inhibitor SW033291

**DOI:** 10.1101/2021.08.15.456403

**Authors:** Alan B. Dogan, Nathan A. Rohner, Julianne N.P. Smith, Jessica A. Kilgore, Noelle S. Williams, Sanford D. Markowitz, Horst A. von Recum, Amar B. Desai

**Author notes:** Denotes equal contribution (Horst A. von Recum, Amar B. Desai). Correspondence; Case Western Reserve University.

## Abstract

As the prevalence of age-related fibrotic diseases continues to increase, novel antifibrotic therapies are emerging to address clinical needs. However, many novel therapeutics for managing chronic fibrosis are small-molecule drugs that require frequent dosing to attain effective concentrations. While bolus parenteral administrations have become standard clinical practice, an extended delivery platform would achieve steady state concentrations over a longer time period with fewer administrations. This study lays the foundation for the development of a sustained release platform for the delivery of (+)SW033291, a potent, small-molecule inhibitor of the 15-hydroxyprostaglandin dehydrogenase (15-PGDH) enzyme, which has previously demonstrated efficacy in a murine model of pulmonary fibrosis. Herein, we leverage fine-tuned cyclodextrin microparticles – specifically β-CD microparticles (β-CD MPs) – to extend the delivery of 15-PGDH inhibitor, (+)SW033291, to over one week.

**Graphical Abstract:** 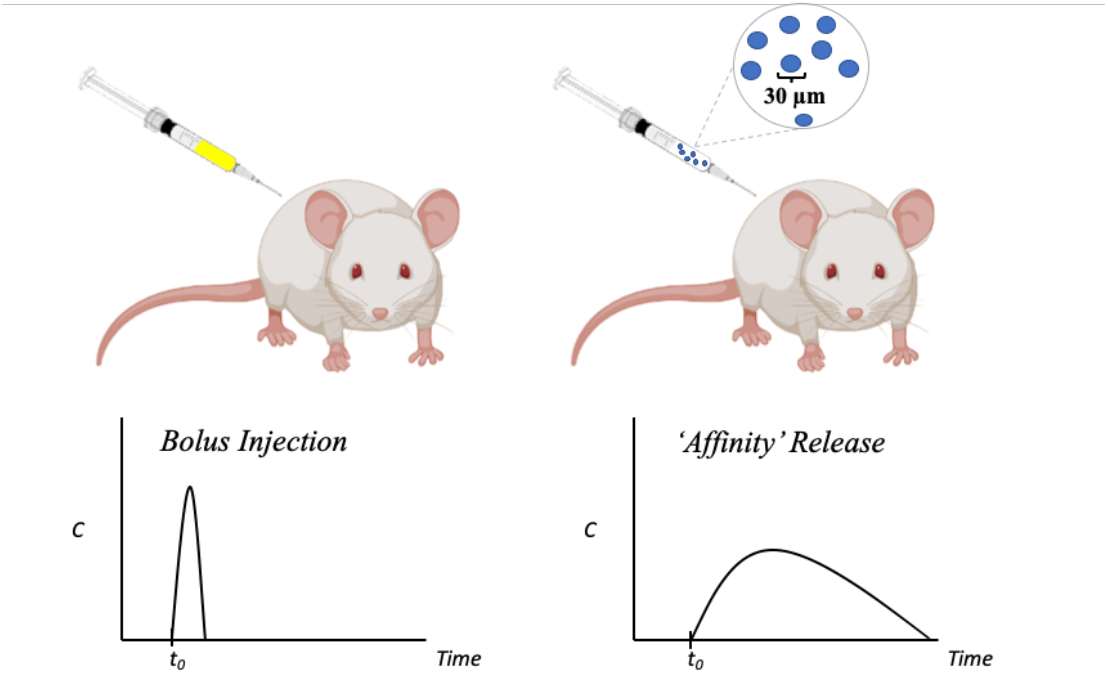

## 1. Introduction

In a normal wound healing response, fibroblast activity is essential for remodeling the extracellular matrix (ECM) after injury. However, in instances of unbalanced or uncontrolled tissue remodeling, excessive deposition of ECM components results in fibrosis, which is a common pathological outcome of several chronic inflammatory diseases. During fibrosis, connective tissue replaces parenchymal tissue, rendering the inflicted tissue partially or completely inflamed or damaged. Clinically, the most common manifestations of fibrotic damage can be seen in chronic diseases such as end-stage liver failure, kidney disease, heart failure, and idiopathic pulmonary fibrosis (IPF)^1^.

Markedly, IPF, a chronic interstitial lung disease characterized by dyspnea, cough and increasing immobility, is particularly prevalent in 60-75 year old patients with a history of smoking and/or occupational exposure to inhaled hazards^2^. While the disease is expected to increase in prevalence in an increasingly aging population, current treatment options for IPF only extend average life to 9-11 years after diagnosis^3^. To date, two drugs have been clinically approved for IPF – pirfenidone and nintedanib – however both small-molecule drugs suffer from short half-lives, adverse drug reactions, high costs, and frequent oral dosing (TID, 3× daily and BID, 2× daily, respectively)^4,5^. Additionally, a large percentage of elderly patients discontinue taking these drugs due to gastrointestinal and other off-target complications^6^.

Inhibition of 15-PGDH, the primary enzyme responsible for Prostaglandin E2 metabolism, using the small molecule (+)SW033291 has recently demonstrated preclinical efficacy in a murine model of bleomycin induced IPF by limiting systemic inflammatory load and reducing pulmonary collagen deposition^7,8^. Further preclinical optimization would involve ensuring that (+)SW033291 can be effectively delivered to the region of treatment with minimized off-target action. Therefore, we propose to expand the clinical utility of 15-PGDH inhibition (PGDHi) by developing a delivery system to administer sustained PGDHi in chronic disease like IPF. We predict that sustained 15-PGDH inhibition via injectable polymer microparticles will be a novel tolerated strategy to not only reduce fibrotic deposition and improve morbidity and mortality in murine pulmonary fibrosis but also reduce the demands on the patient to comply with therapeutic dosing regimens, with clinical implications for a number of additional fibrotic conditions.

To this end, we propose the use of polymerized cyclodextrin microparticles (CD MPs) as a vehicle for the sustained release of the 15-PGDH inhibitor (+)SW033291. CD systems have been previously shown to enhance the solubility and bioavailability of drugs by complexing small-molecule, hydrophobic payloads within its interior pocket, forming an ‘inclusion complex’^9^. Furthermore, CD systems with high densities of neighboring inclusion complexes are able to leverage their thermodynamic interactions with the payload to yield a controlled-release delivery system that can deliver drug for 28-70 days^10–14^.

We believe that based on its molecular structure, SW033291 would be a compatible drug for CD platforms, that the combination device (CD MPs, loaded with SW033291) will require less drug to achieve minimally effective concentrations and will reduce off-target effects in at-risk patient populations, ultimately improving patient outcomes in IPF. Herein, we investigate and characterize the synthesis, loading, *in vitro* kinetics, and preliminary *in vivo* tolerance of CD MPs loaded with SW033291 in murine models to work towards a novel, sustained 15-PGDH inhibitor delivery system.

## 2. Materials and Methods

### 2.1 Materials

β-Cyclodextrin (β-CD) prepolymer, lightly cross-linked with epichlorohydrin was purchased from CycloLab (Budapest, Hungary). Ethylene glycol diglycidyl ether (EGDE) crosslinker was purchased from Polysciences, Inc. (Warrington, PA). (+)SW033291 was provided by Dr. Sanford Markowitz (Case Western Reserve University). The Lad2 cell line was provided by Dr. Dean Metcalfe (NIAID). Phosphate Buffered Saline (1×PBS) was purchased from Millipore Sigma (Burlington, MA). Corning Transwell Plates and all other reagents, solvents, and chemicals were purchased from Thermo Fisher Scientific (Hampton, NH) in the highest grade available.

### 2.2 *In silico* “Affinity” Predictions: cyclodextrin and SW033291

SW033291 was determined to be an appropriate fit for pCD delivery platforms through in silico analysis utilizing both a molecular docking software PyRx, running the Autodock Vina algorithm (Molecular Graphics Laboratory, The Scripps Research Institute, La Jolla, CA) and a cyclodextrin affinity predictor algorithm, developed in our group^15^. Molecular structure data files for SW033291 and cyclodextrin variants (α-CD, β-CD, and γ-CD) were downloaded from PubChem.

### 2.3 Cyclodextrin microparticle synthesis and characterization

Similar to previously established protocols for synthesizing β-CD microparticles (β-CD MPs), epichlorohydrin-crosslinked β-cyclodextrin prepolymer was solubilized in 0.2 M potassium hydroxide (25% w/v) and heated to 60 °C for 10 min^12,16–18^. Light mineral oil was warmed in a beaker with a Tween85/Span85 solution (24%/76%) and stirred at 500 rpm. Ethylene glycol diglycidyl ether (EDGE) was added dropwise, and the solution was vortexed for 2 min before adding the crosslinking solution to the oil/Span85/Tween85 mixture. During crosslinking (3 hours), temperature and mixing speed were maintained at either ‘Method A’: 70 °C at 650 rpms or ‘Method B’: 60 °C at 1500 rpms.

The microparticles were then centrifuged at 200 × g to be separated from the oil mixture and washed with excess hexanes twice, excess acetone twice, and finally deionized water (diH2O) twice. The microparticles were resuspended in diH_2_O, frozen, and lyophilized for 72 hours.

Particle size was determined by a Nikon Eclipse TE300 inverted microscope (Nikon Inc., Tokyo, Japan) and analyzed for particle diameter in ImageJ. Previously studies have cited particle sizes of 81.88 ± 36.86 um with ‘Method A’ synthesis^17^.

Some particle groups were also crushed with a mortar and pestle for 2-3 minutes – referred to as ‘crushed’ - which is intended to reduce particle size after synthesis by ∼70-80 percent^17^.

### 2.4 *In vitro* reduction of Lad2 generated 15-PGDH due to SW033291 loaded β-CD MPs

*In vitro* activity of SW033291 loaded in β-CD MPs was assessed by treating the Lad2 cell line and assessing enzyme inhibition activity, in an assay previously described^19,20^. Lad2 cells have previously shown to highly express 15-hydroxyprostaglandin dehydrogenase (15-PGDH), and therefore were chosen as a model cell-line for assessing PGDHi^19,21^. Briefly, either 1 mL of 600 μM SW033291 (25% DMSO in 2% FBS media) or 20 mg/mL loaded β-CD MPs (2% FBS media) were added to the top portion of the 0.4 μm pore Transwell plate. The bottoms of the plate were seeded with 200k/mL LAD2 cells in 2 mL 2% FBS media and the plates were incubated at 37 °C in static conditions. At each time point, 1 mL of media was sampled from the bottom wells, 0.1 mL of solution was sampled from the top well and replaced with fresh media, and the top wells were transferred to a fresh plate. Enzyme activity from bottom media samples was recorded as counts per minute (CPM) over one hour and normalized to protein input mass (mg).

For each well, 20 mg/mL β-CD MPs were loaded for 24 hours in 250 uM SW033291, washed twice with diH_2_O, and resuspended in media.

### 2.5 Liquid Chromatography/Mass Spectroscopy procedure for SW033291-loaded β-CD MPs analysis

SW033291 concentrations in sampled aliquots were determined via LC-MS/MS in the Preclinical Pharmacology Core at UT Southwestern. 0.1 mL of each sample was incubated at room temperature for 10 minutes with a two-fold volume of acetonitrile, containing 0.1% formic acid, and 50ng/mL tolbutamide to induce solution crashing. Once precipitation occurred, samples were spun at 16,100 × g for 5 min and the supernatant was transferred to a new tube and spun again. Supernatant was transferred to HPLC vials with inserts and analyzed by LC-MS/MS (Sciex 3200 QTRAP mass spectrometer coupled to a Shimadzu Prominence LC). An Agilent C18 XDB column (5 micron packing 50 × 4.6 mm) at 1.5 mL/min with solvents A) Water + 0.1% formic acid and B) MeOH + 0.1% formic acid was used for chromatography prior to introduction of the sample into the mass spectrometer. Concentrations of SW033291 were determined by comparison to a standard curve made by spiking blank matrix with varying concentrations of SW033291 which were processed as for samples.

### 2.6 Improvement of β-CD MPs loading protocol with SW033291

Traditional drug loading procedure (single solvent, 250 uM drug, 24 hour incubation) was found to only fill approximately 20-30% of available host CD molecules within the pCD MPs. To help drive thermodynamic loading of SW033291 into β-CD MPs, 20 mg of dried MPs were mixed with 800 uL of 20 mg/mL SW033291 in DMSO for 1.5 hours, followed by the addition of 200 uL diH_2_O mixed for 0.5 hours. The addition of water is designed to drive the hydrophobic SW033291 into the polymer. The final 4:1 v/v organic:polar mixtures (final concentration of 16 mg/mL SW033291) were then incubated for at least 48 hours at 4°C. Polymers were spun down at 10,000 rpms for 3 minutes and washed twice with 800 uL media/PBS (**Figure 1**).

**Figure 1:**
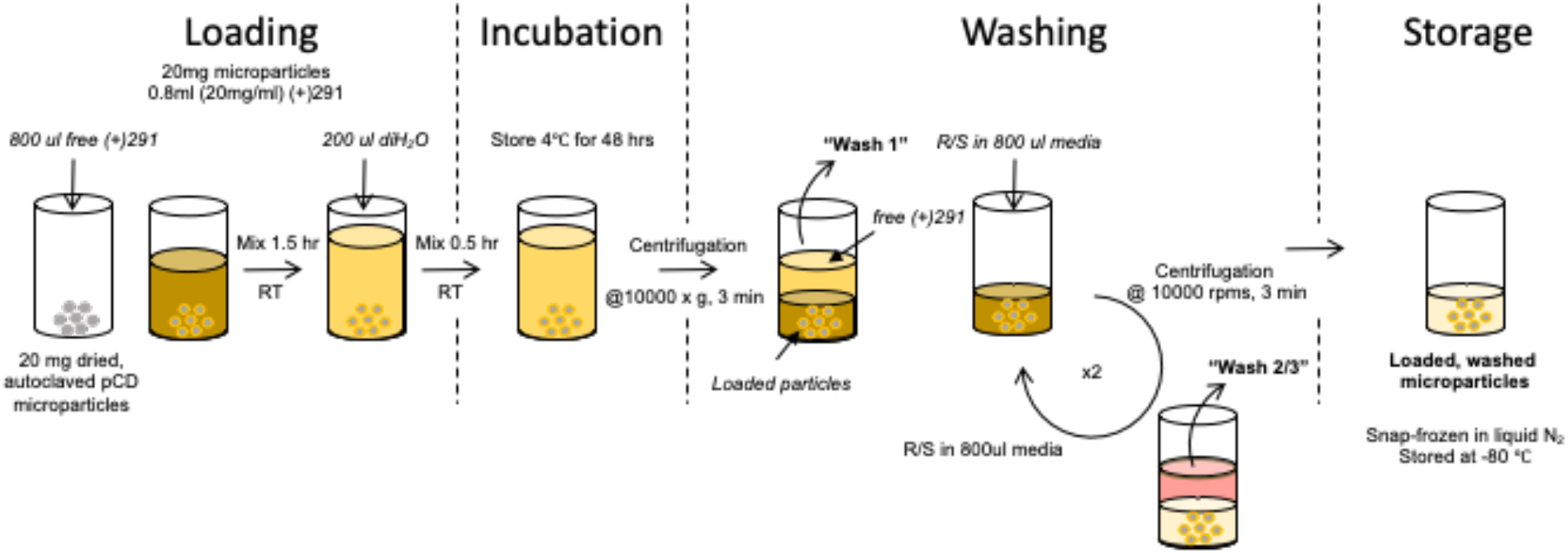
Loading protocol schematic for creating SW033291-loaded β-CD MPs. Prolonging loading to 72 hours helps ensure complexation of SW033291 within cyclodextrin hydrophobic ‘pocket’.

### 2.7 Assessing β-CD MP polymer injectability

Larger diameter β-CD MPs have been previously shown to clog smaller-diameter needles. To ensure our synthesized MPs were injectable from a 29-gauge needle (BD, SafetyGlide Insulin Syringe), both approximate settling time (seconds) and subjective injectability (binary: ‘yes’ or ‘no’) was obtained for β-CD MP formulations.

### 2.8 *In vivo* tolerance murine model

A murine model was used to test β-CD MP tolerance and quantify the extended delivery of SW033291 *in vivo*. 8-week-old female C57/Bl6 mice were purchased from the Jackson lab (CWRU). SW033291-loaded β-CD MPs (2 mg, in 200 uL PBS) were administered via retro-orbital (RO) infusion and peripheral blood was collected into Microtainer serum-separator tubes (Becton-Dickinson) by submandibular cheek puncture at designated time points (n=4 mice per arm; 24, 72, 168, and 336 hrs. post administration). Whole blood was allowed to clot at room temperature and then spun at 6000 × g for 3 minutes to separate the serum. Serum was removed and stored at −80°C prior to sending to the Preclinical Pharmacology Core at UTSW for LC-MS/MS analysis. Analysis was conducted as described above except that the matrix utilized was mouse serum rather than culture media.

### 2.9 Statistical Analysis

Experiments were all carried out in triplicates, unless otherwise stated and data is represented as mean ± standard error. *In vivo* data represents the average of n=4 mice and error bars also represent standard error of the mean.

## 3. Results

### 3.1 SW033291 is predicted to have significant affinity for β-CD host *in silico*

Before beginning *in vitro* studies, we confirmed *in silico* that a cyclodextrin delivery platform would be compatible with (+)SW033291’s chemical structure. Utilizing two methods of affinity prediction - docking simulations and quantitative structure activity relationship (QSAR) - we found that SW033291 had the highest binding affinity with the β form of cyclodextrin (**Figure 2a**). Docking simulations also revealed that the central moiety that binds with cyclodextrin includes SW033291’s central heterocyclic rings and sulfoxide group (**Figure 2b**). Based on previous studies, we predicted a binding affinity of −23 KJ*mol^−1^ would result in a window of release of about 1-2 weeks^16,22–24^.

**Figure 2:**
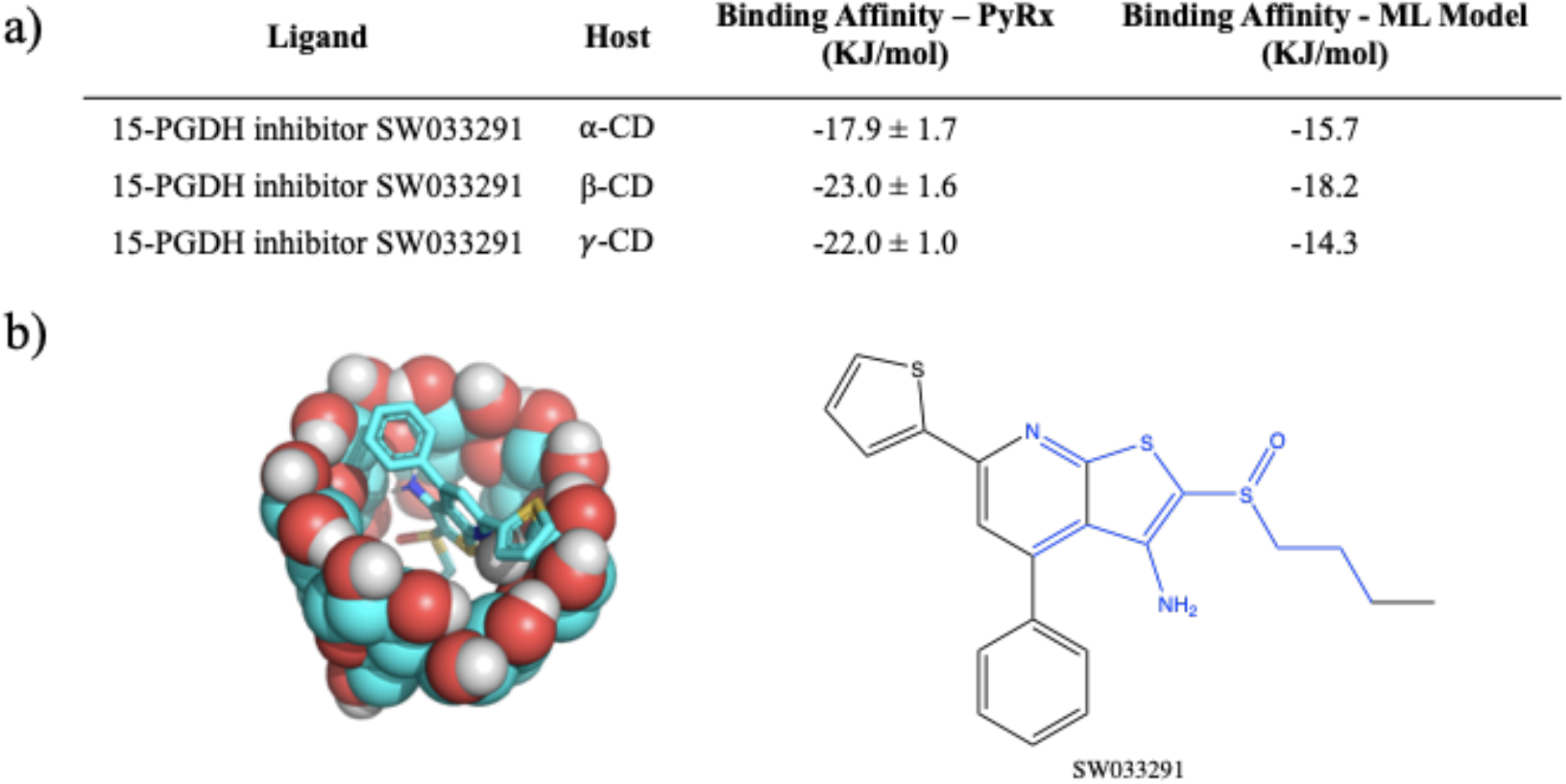
**a)** Affinity binding simulations of 15-PGDH inhibitor SW033291 (CID:337839) complexation with α, β, and γ cyclodextrin (CD) in both PyRx and a machine learning algorithm for affinity prediction^15^. **b)** Molecular structure *in-silico* model demonstrating inclusion complex formation between small molecule drug SW033291 binding to the inner pocket of β-cyclodextrin. Regions in blue are complexed within the cyclodextrin ‘pocket’.

### 3.2 Lowering temperature of polymerization and increasing speed of mixing generates smaller β-CD MPs

Established protocols for synthesizing β-CD MPs generate diameters that were not suitable for small-gauge needle injections, therefore, we sought to generate MPs with smaller diameters to allow for injectability. To this end, we modulated parameters during polymerization - specifically temperature and mixing speed during polymerization - to generate smaller microparticles. We also explored manually crushing MPs with a mortar and pestle, which was previously reported to reduce particle diameters by 30%^17^. By reducing temperature and raising mixing speed to 1500 rpm, we generated MPs ∼40 um in diameter, a 50% reduction from the original protocol (**Figure 3**). Microparticles were then loaded for 24 hrs. with (+)SW033291, and final loading concentrations were reported after drug leaching.

**Figure 3:**
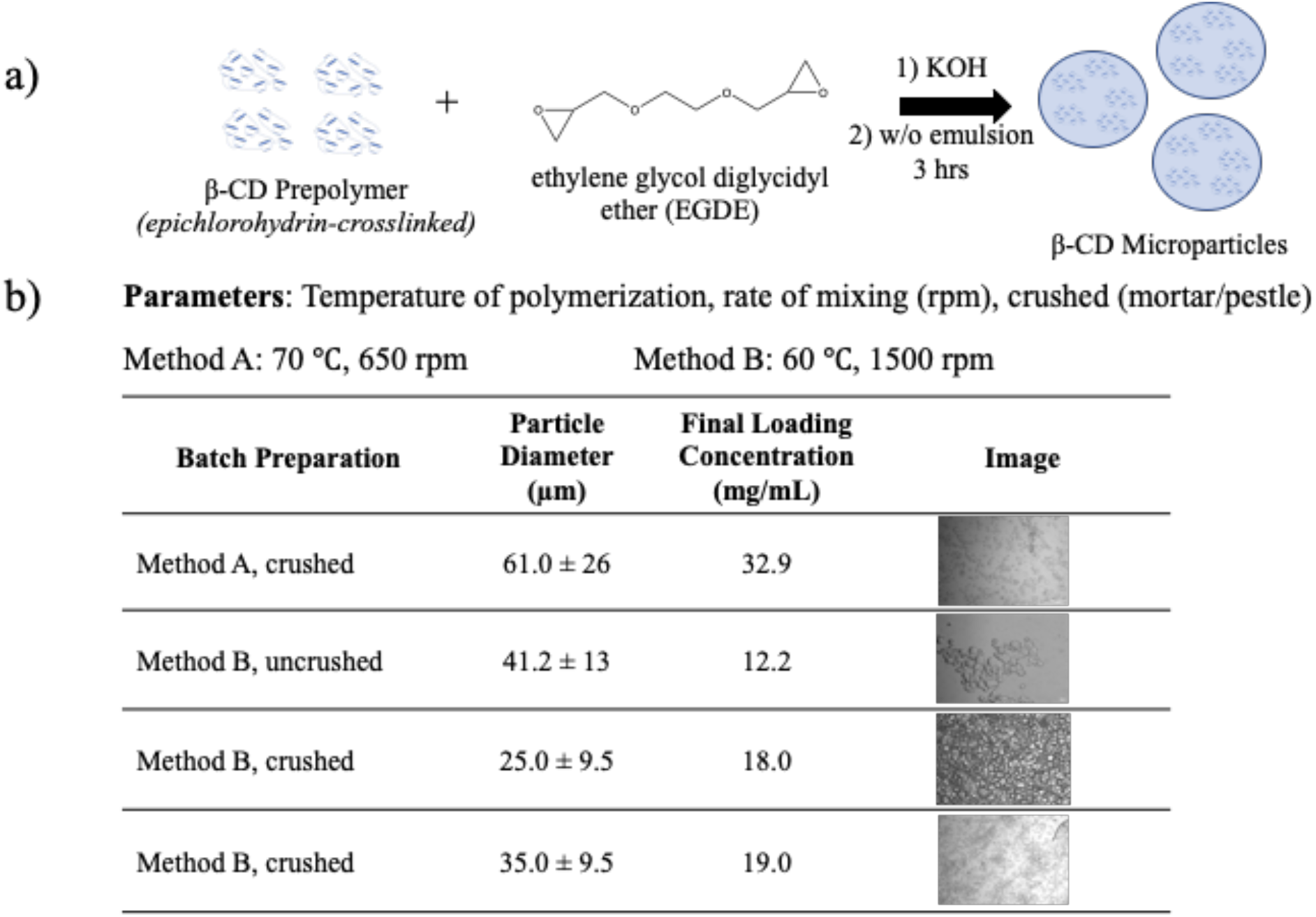
**a)** Overview of β-CD microparticle preparation. β-CD prepolymer is polymerized with EDGE for 3 hours at elevated temperature (60-70 °C) and formed in a water/oil (w/o) emulsion. **b)** Microparticle batches synthesized with varying temperature and mixing speeds. Particles were then loaded with (+)SW033291 for 24 hours (DMSO). Particle diameters were determined by ImageJ, and represented as the mean ± S.D. Final loading concentrations were determined via LC/MS at UTSW

### 3.3 SW033291 released from β-CD MPs has prolonged Lad2 inactivation compared to free drug

To confirm drug activity after delivery from β-CD MPs, (+)SW033291 activity was analyzed *in vitro* by proxy of analyzing supra-physiological concentrations of Lad2 enzyme activity inhibition. Using Method A, uncrushed particles (final loading concentration: 11.9 mg/mL), we found that β-CD MP - released (+)SW033291 was effective at reducing LAD2 enzyme activity comparable to 5 uM free drug after 72 hours (**Figure 4**). LAD2 controls and LAD2 treated with unloaded β-CD MP controls saw unremarkable changes in enzyme activity, confirming that β-CD alone does not contribute to enzyme inactivation. Drug release kinetics were also observed to exhibit increasingly zero-order kinetics compared to bolus administration, confirming β-CD MPs slowed the rate of release of (+)SW033291 over a three-day period (**Figure S2**).

**Figure 4:**
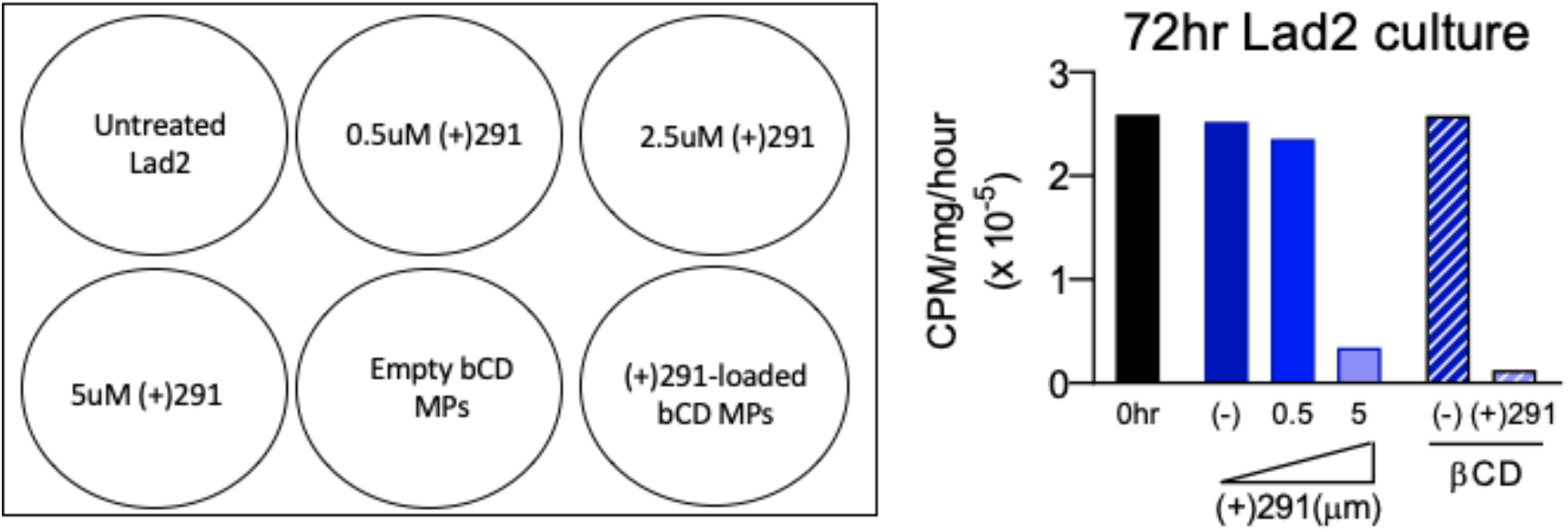
Incubation study of (+)SW033291-loaded β-CD MPs with Lad2 cells indicated sustained delivery in co-culture inhibits enzyme activity 72 hrs post-administration with similar efficacy as bolus SW033291 administration.

### 3.4 β-CD MP Loading Optimization

To help address the relatively low observed ‘final loading concentration’ seen in the 24-hour loaded β-CD MP, we altered the loading protocol to encourage increased packing of drug within the polymer particles. Comparing ‘24-hour loading’, which used 24-hour incubation in (+)SW033291 dissolved in DMSO, with ‘72-hour loading’, which is previously described as mixing a polar and nonpolar solvent to encourage complexation of the hydrophobic drug within the CD pockets. We found that altering this procedure yielded final loading concentrations 4 to 14-fold greater compared to our previous protocol (**Figure 5**).

**Figure 5:**
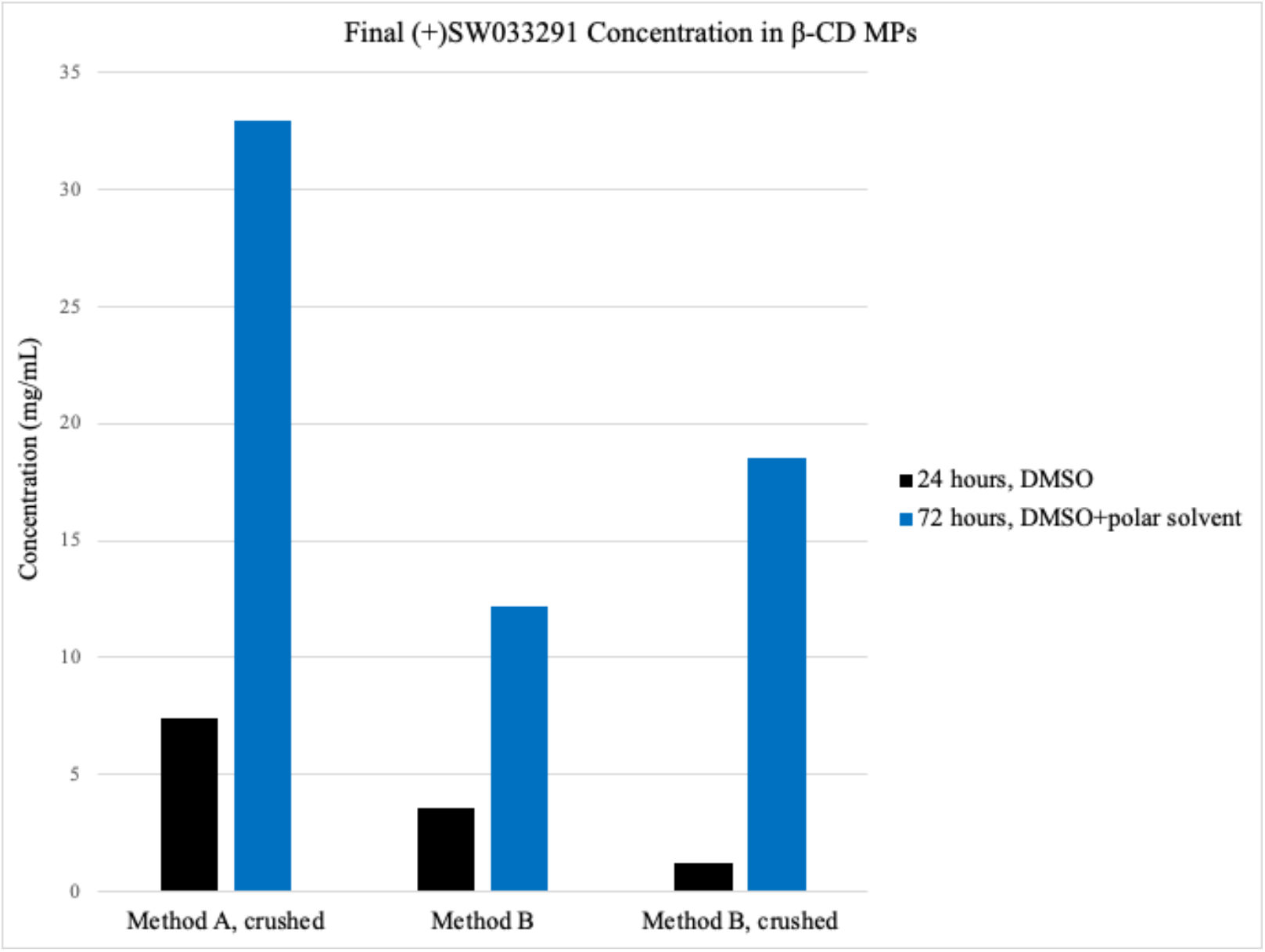
Final loading concentrations for β-CD MP formulations utilizing a ‘24-hour’ loading protocol and our investigated ‘72-hour’ loading protocol, which also uses a polar solvent (water) to help drive complexation.

### 3.5 At lower concentrations, β-CD MPs are injectable but are prone to clumping

Before moving to an *in vivo* model, we aimed to investigate if smaller-diameter β-CD MPs are suitable for injection from 29-Gauge syringes. In a serial dilution of 40 mg/mL, which was determined to be the maximum concentration of MPs achievable in an aqueous medium without immediate crashing, we found that the formulations were injectable ≤10 mg/mL, however issues with clumping were observed (**Table 1**). At a lower concentration of ≤1.3 mg/mL there was notedly less clotting observed. Moving forward *in vivo* we selected 10 mg/mL in order to ensure maximum β-CD MP delivery in a suitable volume (∼200 uL).

**Table 1:** Serial dilution of β-CD MPs in 1× PBS. Approximate settling times were measured subjectively when depositions of polymer were first observed to crash out from solution. ‘Injectability’ was rated as a binary ‘yes’ or ‘no’ qualitative observation.

| Concentration of β-CD<br>MPs (mg/mL, d=35 μm) | Approximate<br>Settling Time<br>(s) | Injectable?<br>(29 G needle) |
| --- | --- | --- |
| 40 | 5 | no |
| 20 | 9 | no |
| 10 | 16.6 | yes |
| 5.2 | 20 | yes |
| 2.6 | 23 | yes |
| 1.3 | 30 | yes |
| 0.7 | >60 | yes |

### 3.6 (+)SW033291 loaded β-CD MPs prolonged payload delivery over 1 week and were well tolerated in murine model

SW033291-loaded β-CD MPs (2 mg, in 200 uL PBS) were administered via retro-orbital (RO) infusion to 8-week-0ld female C57/Bl6 mice (80mg/kg) and peripheral blood was collected into Microtainer serum-separator tubes (Becton-Dickinson) by submandibular cheek puncture at designated time points (n=4 mice per arm; 24, 72, 168, and 336 hrs. post administration). Within the cohort of 16 mice, we observed no adverse events following RO infusion within the 2-week study. Serum drug concentrations (n=4 at each timepoint) revealed that (+)SW033291 was still present after 1-week post administration of SW033291 loaded β-CD MPs, eventually becoming undetectable in three out of four mice at the 2-week timepoint (**Figure 6**).

**Figure 6:**
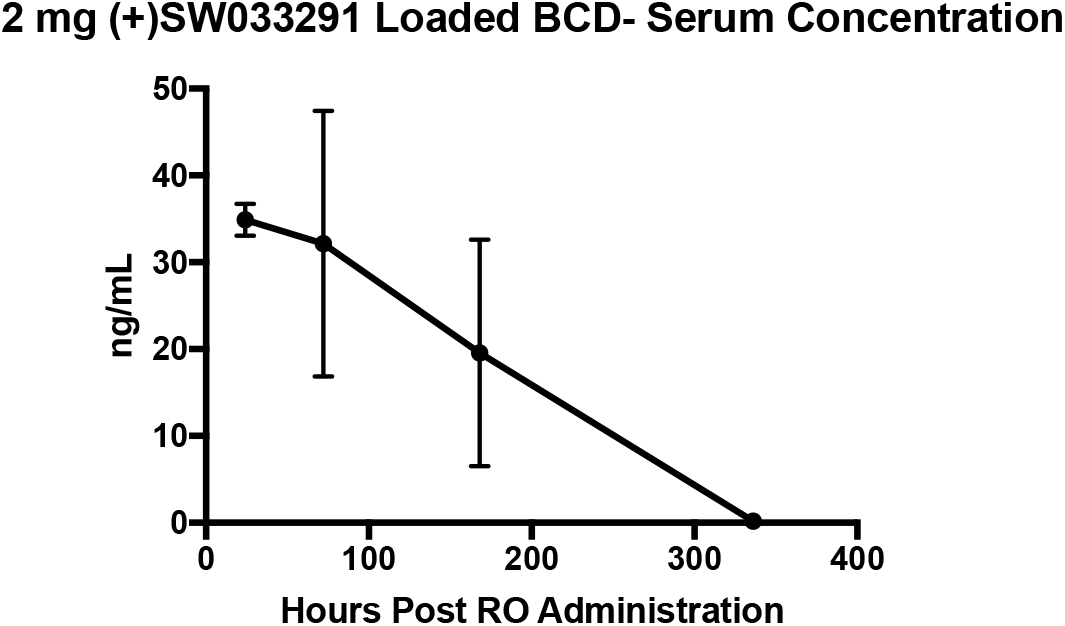
Serum (+)SW033291 concentrations collected from mice (n=4 at each timepoint) following RO infusion of (+)SW033291 loaded β-CD MPs. Error bars are representative of the standard error of the mean.

## 4. Discussions

While pCD delivery systems have been used to extend the delivery of other small-molecule therapeutics, this is the first study showing that pCD is a compatible sustained release platform for PGDHi by (+)SW033291. Using *in silico* methods for affinity prediction, we found that SW033291’s structure was most compatible with β-CD (**Figure 2**). We found that (+)SW033291:β-CD interactions had comparable free energy of binding to existing drugs that have been successfully delivered from pCD platforms (e.g. doxorubicin) (**Figure S1**). We then fine-tuned existing synthesis protocols for our lab’s β-CD MPs, generating a smaller-diameter microparticle around 30-35 μm (**Figure 3**), confirming that decreasing temperature and increasing rate of stirring during polymerization in a water/oil emulsion generates smaller pCD particles.

Effective delivery of (+)SW033291 from β-CD MPs was also confirmed, first *in vivo* by effectively reducing supra-physiological enzyme activity in LAD2 cells (**Figure 4**), and secondly *in vivo* in healthy mice (**Figure 6**). Drug kinetics *in vitro* revealed that (+)SW033291 was taking advantage of pCD inclusion, as the time of release was extended up to 6 days - compared to a 1 day bolus input - and exhibited increasingly zero-order kinetics (**Figure S2**).

(+)SW033291 loading protocols for β-CD MPs were also improved after discovering 24-hour incubation times in DMSO were insufficient to promote drug inclusion. To this end, we modified existing loading protocols by extending incubation time to 72 hours and adding an aqueous component to the loading solution to help promote drug inclusion to pCD pockets (**Figure 1, 5**). This further confirms that drug inclusion within cyclodextrin in highly dependent on solvent contact time and respective solubility of the drug within the drug-loading solvent.

While it has been previously reported that pCD polymers are relatively well tolerated *in vivo*, we found that 2 mg of β-CD MPs administered via RO infusion was tolerated without ramifications in all mice within our cohort (**Figure 6**). We also observed that from a single dose of (+)SW033291-loaded β-CD MPs was able to produce a drug delivery profile of over 1 week (**Figure 6**).

This study represents the groundwork for developing an extended drug delivery system for anti-fibrotic therapies. While we were able to access global blood serum levels of (+)SW033291, future studies will focus on analyzing the biodistribution of the β-CD MPs and whether or not drug localization is achieved in regions of interest, specifically in the pulmonary tract and hepatic system^20^. While the window of delivery that was achieved *in vivo* greatly surpassed bolus injection, the amount of drug released from 2 mg most likely would be subtherapeutic. Therefore, we propose to increase the frequency of β-CD MPs administration in future models to help ensure that drug concentrations surpass minimally effective concentrations. We also aim to reduce MP clumping during administration, in which either administration protocols or synthesis quality (reducing polydispersity) can be changed to make MP injections easier^25^. Ultimately, we aim to further optimize the pCD platform for sustained delivery of (+)SW033291 and assess the efficacy of sustained delivery PGDHi as a therapeutic strategy in multiple models of chronic fibrotic disease.

## 5. Conclusion

Herein, we present the foundation for creating an injectable, polymer microparticles system for sustained 15-PGDH inhibition. As age-related fibrosis increases in prevalence, long-term delivery of antifibrotic therapeutics will greatly improve upon current standards of care and augment the clinical usability of potent small-molecule fibrosis inhibitors, like (+)SW033291. Extending delivery of (+)SW033291 for over a week *in vivo* presents the first step in creating a novel tolerated strategy to reduce fibrotic deposition and improve morbidity and mortality in murine pulmonary and hepatic fibrosis. Future success in (+)SW033291 prolonged delivery would also have additional clinical implications for a number of fibrotic conditions.

## Author Contributions

H.A.R. and Amar B.D. devised the project. Alan B.D. developed the technical procedures and performed the *in vitro* experiments under the supervision of H.A.R. and Amar B.D. In vivo work was completed by Amar B.D. J.A.K. evaluated concentrations of (+)SW033291 for both *in vitro* and *in vivo* analyses under the supervision of N.S.W. Under the supervision of H.A.R and Amar B.D., Alan B.D. wrote the manuscript; all authors read or edited the manuscript.

## Acknowledgements

This work was supported by NIH grants 5R00HL135740-04, R35 CA197442, and by the Radiation Resources Core Facility (P30CA043703) of Case Western Reserve University, as well as the Preclinical Pharmacology Core of UT Southwestern.

We also thank Dr. Joseph Ready of UT Southwestern for performing QC on (+)SW033291 and providing additional scientific insight.

## Supplemental Figures

**Figure S1:**
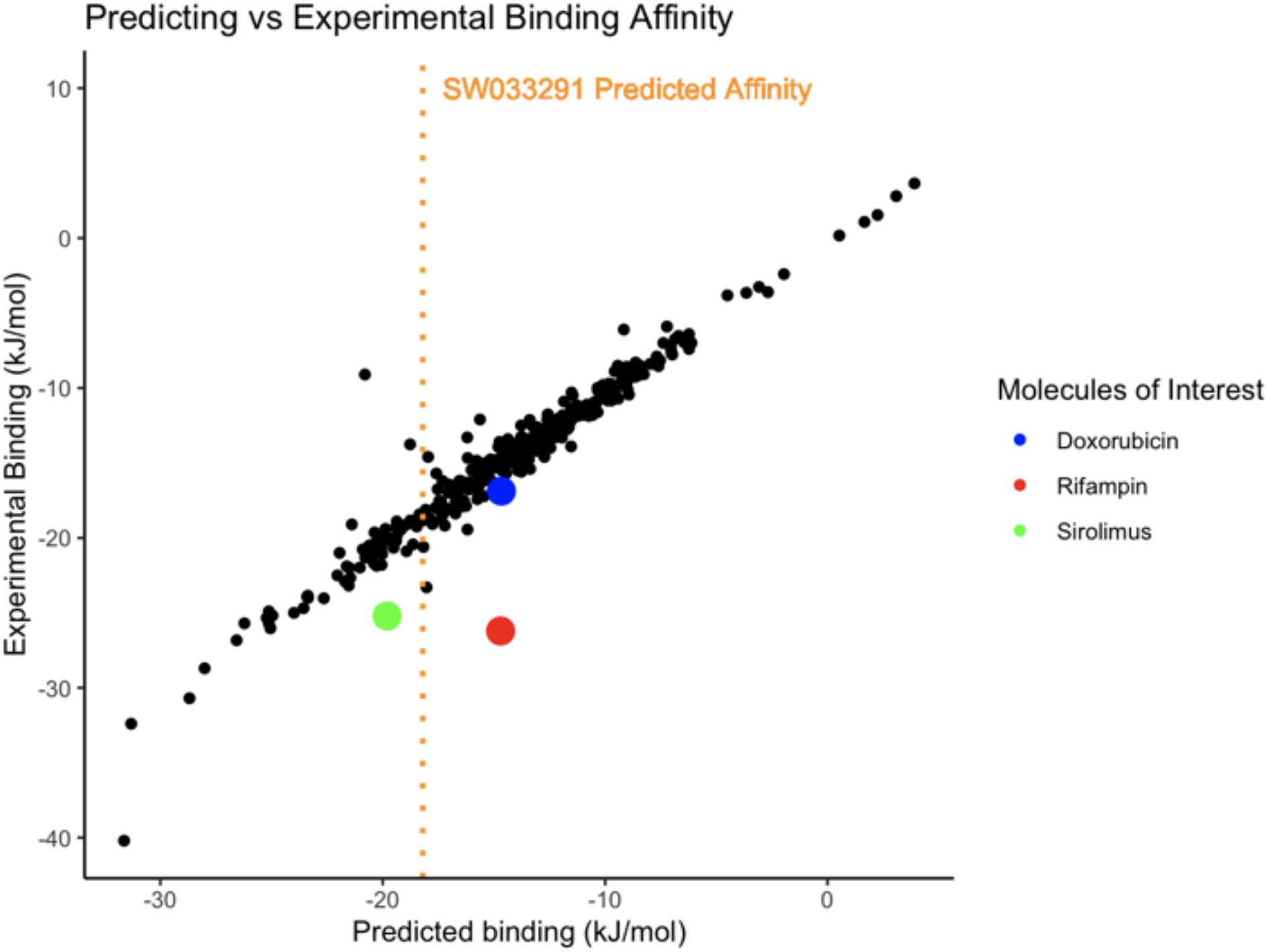
Affinity prediction machine learning model with training set (black) and highlighted drugs (“molecules of interest”) that were previously compatible with β-CD platforms. The dotted orange line is representative of SW033291’s predicted affinity to β-CD based on chemical descriptors.

**Figure S2:**
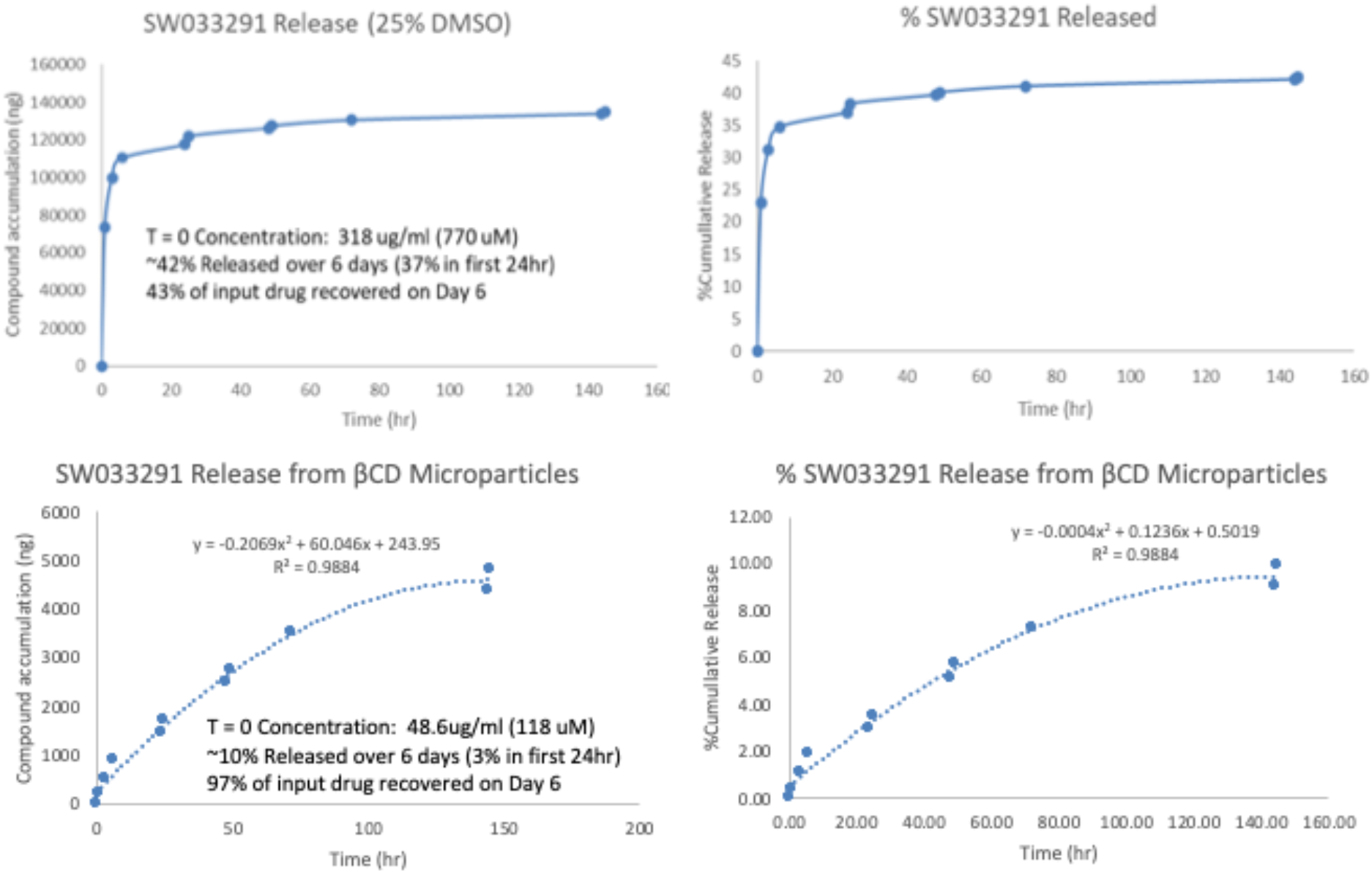
Drug release kinetics sampled from top transwell plates for both bolus administered SW033291 and β-CD MPs loaded with SW033291 (incubated for 24 hours).

